# Development of a Matrix-Matched Calibration Curve for Multi-Site Quantification of Neu5Gc-Bearing N-Glycans

**DOI:** 10.64898/2026.07.14.738351

**Authors:** Nicholas J DeBono, Edward S. X. Moh, Jessica Poole, Nicolle H. Packer, Christopher J Day, Michael P. Jennings, Daniel Kolarich, Christopher Ashwood

## Abstract

*N*-glycolylneuraminic acid (Neu5Gc) has been repeatedly associated with human cancer, but reliable detection has remained elusive, generating controversy regarding its presence in human samples. To address this, matrix-matched calibration curves, which have been pioneered in proteomics and metabolomics for assessing changes in complex mixtures, were measured of released *N*-glycans at four orders of magnitude dynamic range in defined mixtures, systematically benchmarking Neu5Gc-containing *N*-glycan detection across multiple LC-MS platforms and sites. Orthogonally, the gold-standard analytical method, consisting of fluorescence detection of labelled monosaccharides separated by LC, was applied to the same samples, yielding absolute concentrations of Neu5Gc.

LC-MS demonstrated an extended detection range of three or more orders of magnitude while retaining intact *N*-glycan measurement, improving assay specificity and enabling detection of the variety of Neu5Gc-bearing *N*-glycans. By combining orthogonal dimensions of evidence, including chromatographic separation, isotopic distribution matching, and composition-confirming MS/MS, LC-MS confidently resolved Neu5Gc signals from noise, even at low abundance. In comparison, DMB-LC-FLR was limited to two orders of magnitude dynamic range, insufficient for detection of Neu5Gc in commercially available pooled human sera. These findings strongly support that DMB-LC-FLR assay specificity and sensitivity are insufficient for Neu5Gc detection in human samples due to noise overwhelming the Neu5Gc signal. By establishing a reusable benchmarking framework for future glycomic studies, we aim to use LC-MS to improve the measurement of Neu5Gc in clinical samples.

## Introduction

Aberrant glycosylation of cancer tissue is a repeatedly reported difference from healthy human tissue, across multiple cancer types (Meany and Chan 2011, Tuccillo, De Laurentiis et al. 2014, Munkley and Elliott 2016). A key glycosylation change that is consistently found is the presence of *N*-glycolylneuraminic acid (Neu5Gc) (Wang, Shewell et al. 2023). Neu5Gc is traditionally thought not to be expressed in healthy human tissue due to an inactive cytidine monophosphate *N*-acetylneuraminic acid (Neu5Ac) hydroxylase (CMAH) enzyme, responsible for converting Neu5Ac to Neu5Gc (Suzuki 2006). In human cancer tissue and secretion material, Neu5Gc has been reported at very low levels, particularly when compared to the more common sialic acid, Neu5Ac (Shewell, Wang et al. 2018, Shewell, Day et al. 2022, Shewell, Day et al. 2023, Wang, Shewell et al. 2023, De Bisscop, Shewell et al. 2026). Despite the clear association of Neu5Gc with cancer tissue, and the serum of people with cancer, the very low abundance of Neu5Gc has hampered progress towards developing a specific and accurate biomarker, with a specific Neu5Gc lectin only having been developed recently (Wang, Shewell et al. 2018). While lectin specificity can at times be unparalleled, more direct evidence such as that provided by mass spectrometry (MS) is often required for further confirmation.

In part, the lack of development of a Neu5Gc related disease biomarker reflects the limited sensitivity in available instrumentation to detect the presence of lowly abundant Neu5Gc in human samples. Determination of relative limits of detection for lowly abundant molecules can have direct benefits for downstream biomarker detection and validation, potentially leading to more specific disease diagnosis and treatment (Zhang, Whiteaker et al. 2019). Mass spectrometry-based analytical methods for glycomics can be performed in a multitude of ways, including as glycopeptides or glycolipids, and as released glycans, both labelled and unlabelled (Fan, Lu et al. 2025). Usually, quantative sialic acid analysis is performed via 1,2-diamino-4,5-methylenedioxybenzene (DMB)-liquid chromatography (LC)-FLR (Sato, Inoue et al. 1999), which involves acid hydrolysis of the glycan to release the sialic acids and subsequent labelling with the fluorescent molecule DMB. DMB-labelled sialic acids are typically analysed using a liquid chromatograph coupled to a fluorescence detector, however this method is also compatible with mass spectrometry detection(Omoto, Yamakawa et al. 2025). Reducing glycans to their monosaccharide constituents is destructive and results in a loss of overall structural information and thus, biological context. In contrast, released glycan analysis by MS has the potential to provide both structural and compositional information on glycans.

MALDI- and LC-MS can measure released, intact glycans, providing insight into how monosaccharides are linked together and reflecting overall biosynthetic pathways. Analysing intact structures can enhance specificity, for example by distinguishing Sialyl Lewis epitopes that contain both fucose and sialic acid linkages (Palmisano, Larsen et al. 2013). While these methods generally provide relatively unbiased profiling of glycans, with some targeted assays reported (Ruhaak, Xu et al. 2018, Ashwood and Cummings 2025), untargeted analyses that quantify tens to hundreds of structures in a sample seldom evaluate the quantitative accuracy of these measurements.

To assess whether changes in glycan abundance measured by MS truly reflect changes in concentration, calibration curves are essential. Quantitative accuracy depends on whether an increase in analyte concentration produces a proportional increase in measured signal, with the lower limit of quantitation (LLOQ) defining the point below which this relationship no longer holds (Currie 1995). In glycomics, the relevant matrix depends on the sample and the glycan release method, and comprises both the glycans of interest and any co-eluting contaminants present during LC-MS injection. Matrix-matched calibration curves (MMCCs) have been widely adopted in other -omics fields to enable accurate, multiplexed quantitation. They are routinely applied in proteomics for thousands of peptides (Pino, Searle et al. 2020) and in metabolomics or environmental analyses for tens to hundreds of analytes (Kang, Hick et al. 2007). This approach has yet to be systematically applied to glycomics, despite its potential to improve quantitative reliability across multiple glycan compositions and structures. Traditional calibration methods remain limited by the availability and cost of isotopically labelled internal standards (e.g. 13C-labelled Neu5Gc glycans), so matrix-matched dilution strategies offer an attractive route to assess quantitative accuracy in complex glycan mixtures across LC-MS platforms.

We hypothesise that there has been an under-reporting of Neu5Gc related glycan structures in MS data from human glycomic samples, due to a combination of restrictive search spaces in data analysis and low levels of abundance of Neu5Gc containing glycans. To assess the detection and quantitation limits of released glycans, with the goal of determining sensitivity towards Neu5Gc in human samples, we provided a set of samples to multiple independent MS labs with well-established routine glycomics workflows. These samples contained initially evaluated matrix matched standards of reduced *N*-glycan alditols carrying Neu5Gc containing glycans in a serial dilution. Appropriate analysis of these standards allows for a matrix matched calibration curve to be developed (Pino, Searle et al. 2020), determining limits of detection (LOD) and LLOQ for Neu5Gc containing *N*-glycans against a complex background. To the best of our knowledge, this is the first time this has been applied to analyses of released *N*-glycans. Conclusions drawn from this study provide insights into instrumentation-specific sensitivity limits, contextualizing platform-specific findings. A comparative study was also performed against the traditional labelled DMB-LC-FLR analysis, providing information on differing levels of sensitivity with a matched sample set.

## Results

### Experimental design

*N*-glycans enriched in Neu5Gc were released and profiled from a proprietary source. These *N*-glycans were predominantly Neu5Gc-modified, and were primarily biantennary, complex class (**Figure 1A**). Due to these characteristics, this sample was suitable as the Neu5Gc target glycome for the MMCC (**Figure 1B**). As MMCCs require a require a closely matched matrix that is free of the target analytes, human serum *N*-glycans were chosen as a matrix because they are structurally similar (complex, biantennary) and contain only trace (0.01%) amounts of Neu5Gc (**Figure 1C**).

**Figure 1.**
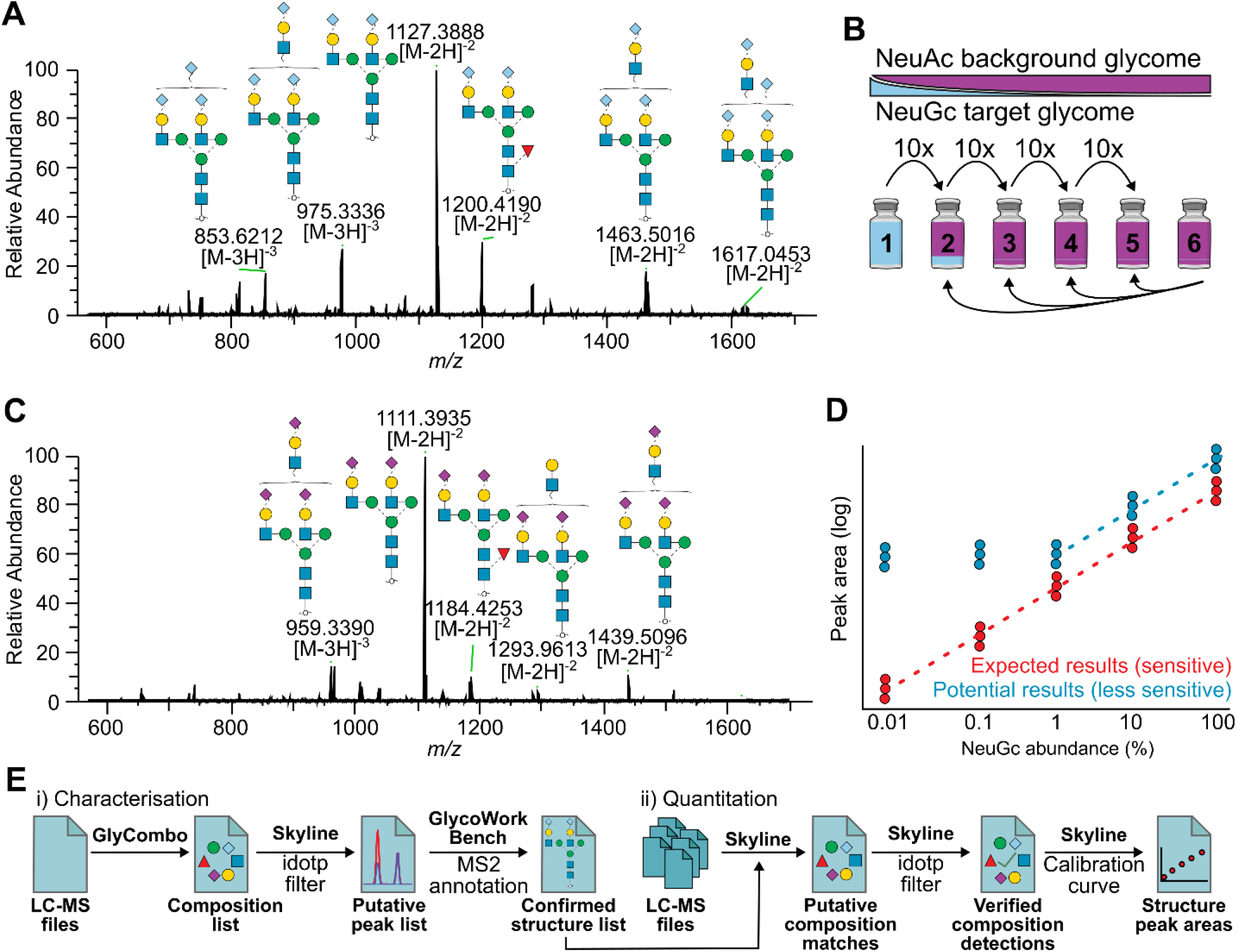
Development and application of a matrix-matched calibration curve to quantify Neu5Gc-bearing *N*-glycans. **A** Neu5Gc enriched sample *N*-glycan profile (Tube 1). **B** Design of the Neu5Gc benchmark set. **C** Human serum *N*-glycan profile to be used as the background matrix. **D** Expected results when analysing the Neu5Gc benchmark set. **E** Data analysis workflow for the multi-site study.

To develop a set of samples for benchmarking Neu5Gc detection across multiple sites, a total of six samples were generated by serially diluting the Neu5Gc target *N*-glycome with the Neu5Ac background *N*-glycome, covering four orders of magnitude dynamic range (**Figure 1D**). A multi-site laboratory study was initiated in which each set of six tubes (the benchmark set), prepared at Site 1, was analysed. Each participating group was sent at least one benchmark set for oligosaccharide analysis by LC-MS and/or monosaccharide analysis by LC-fluorescence (FLR).

Data were collected from each site (for a total of 4 LC-MS instruments) and subjected to a data analysis workflow (**Figure 1E**) to obtain qualitative and quantitative values for each glycan observed. To build the glycan list for assessing identification and quantitation quality at each Neu5Gc concentration, GlyCombo (Maia Kelly 2024) was used to identify glycans subjected to MS2, with Skyline (MacLean, Tomazela et al. 2010, Adams, Pratt et al. 2020) subsequently used to filter for high quality glycan identifications with isotopic distributions matching expected. Structure was then verified by MS2 annotation in GlycoWorkbench (Ceroni, Maass et al. 2008). This list of glycans was then used as the target list within Skyline, with isotopic distribution filtering only performed to maintain quality before extracting glycan peak areas.

### Qualitative comparisons for Neu5Gc detection

Neu5Gc glycans can be distinguished from their Neu5Ac counterparts using several independent measurements across the PGC-LC-MS workflow. At the MS1 level, in addition to the difference in mass (291.10 vs 307.09 Da), both the monoisotopic precursor mass and the approximate ratio of the naturally occurring isotopes (^15^N, ^13^C, ^2^H, ^18^O) can be predicted, enabling comparison between the observed isotopic distribution and the theoretical distribution. The idotp metric (with 1 being a perfect match) can thereby confirm that an observed peak corresponds to the expected glycan chemical formula, and can distinguish true glycan signal from chemical and/or detector noise.

Since MS2 is triggered on a single precursor *m/z*, complementing it with the predicted isotopic distribution allows the full MS1 information in each raw file to be assessed, limiting the impact of noise on detection. As expected, idotp values were highest for the Neu5Gc-enriched sample analysed without background matrix and decreased with decreasing concentration, with values across almost all LC-MS platforms inversely correlated with dilution factor (**Figure 2A**).

**Figure 2.**
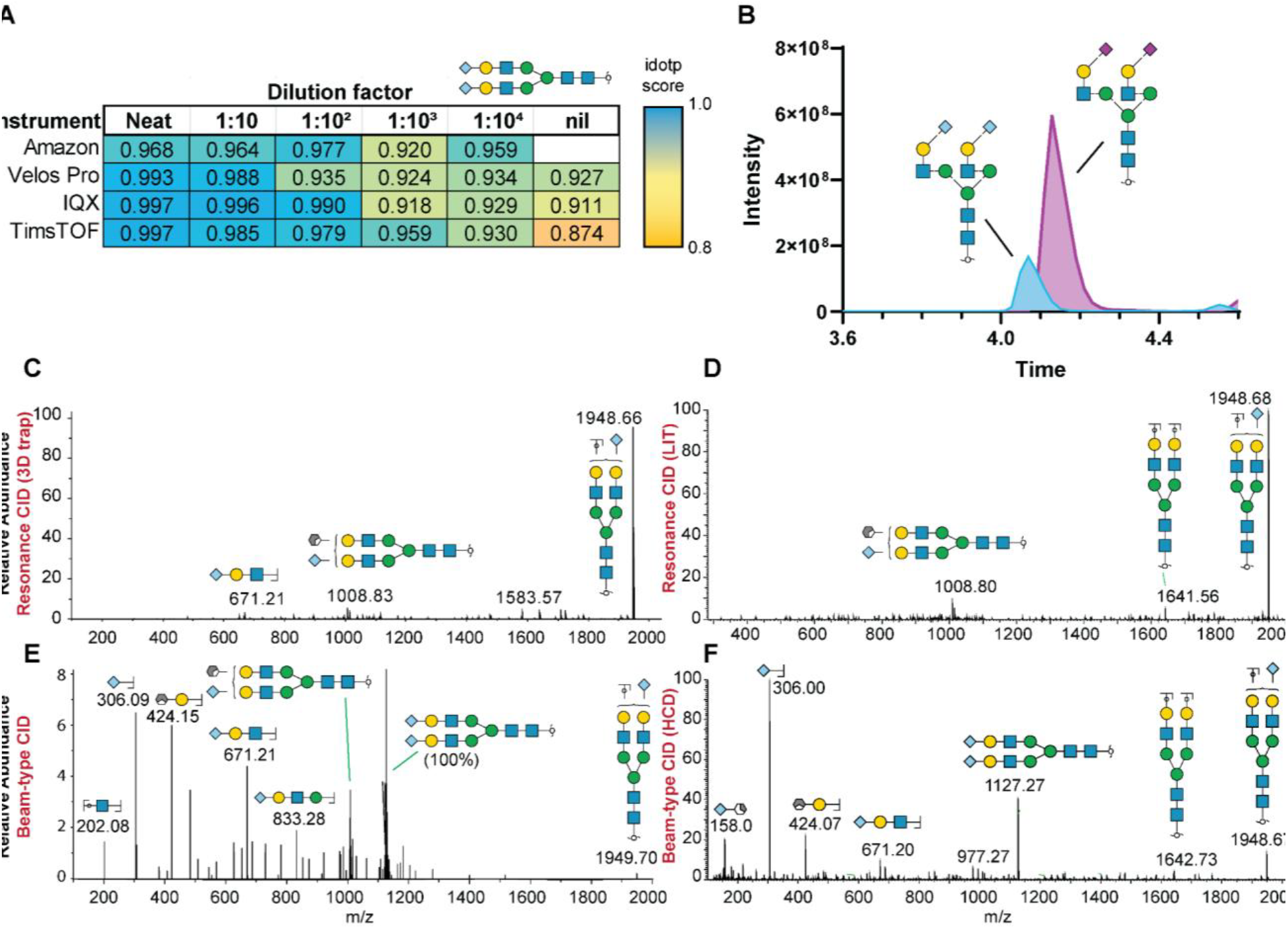
Detection and fragmentation of Neu5Gc biantennary glycan. **A** Average idotp of biantennary Neu5Gc glycan from all participating instruments according to dilution factor. **B** Partial co-elution of Neu5Gc and Neu5Ac variants of the same glycan. Chromatographic trace generated with data from Site 1. Different fragmentation methods from different instruments result in varying fragmentation spectra, as exemplified by resonance 3D trap CID (**C**), linear ion trap resonance CID (**D**), beam-type CID (**E**), and HCD (**F**) data shown.

Chromatographically, Neu5Gc glycans consistently elute ahead of their Neu5Ac counterparts on the PGC, a separation best resolved at a 1:10 dilution factor. This separation matters for targeted analysis, as the two species partially overlap and are therefore prone to mutual interference and ion suppression (**Figure 2B**). Neu5Gc glycans also display a defined ion mobility profile that can be exploited to further separate them from Neu5Ac (**Supplementary Figure 2**).

At the MS2 level, fragmentation behaviour is technique-dependent. Resonance collision induced dissociation (CID), as used on the amaZon^™^ and Velos Pro instruments, produces no detectable Neu5Gc monosaccharide signals due to the 1/3 rule that applies to ion trap instruments(March 1997, Ashwood, Abrahams et al. 2017); one of the smallest product ions observed is 671.22 *m/z* ([M-H]^-^) while charged losses of Neu5Gc from the precursor can also be observed (**Figure 2C–D**). In contrast, beam-type CID generates the Neu5Gc monosaccharide at 306.08 *m/z* as the most abundant product ion, except in the timsTOF Pro data where the unfragmented precursor dominated (**Figure 2E–F**); additional fragments such as Hex+Neu5Gc are informative and detectable across all platforms.

Because the fragmentation pattern depends on the dissociation method, search strategies must be tailored accordingly. Diagnostic-fragment approaches that rely on the Neu5Gc monosaccharide, such as that of Helm *et al*. (Helm, Grünwald-Gruber et al. 2021) are unsuitable for resonant CID, where the low-mass cut-off (approximately one-third of the precursor *m/z*) prevents detection of the monosaccharide, and spectral library-based searching is likewise fragmentation method-specific (Helm, Grünwald-Gruber et al. 2021).

### Quantitative comparisons for Neu5Gc detection

A calibration curve, as illustrated in **Figure 3A**, serves to define the quantifiable range, bounded by the lower (LLOQ) and upper (ULOQ) limits of quantitation, corresponding to noise and saturation respectively. Variation, represented by the difference in intensity between replicates at the same concentration, ideally shows less than 20% coefficient of variation.

**Figure 3.**
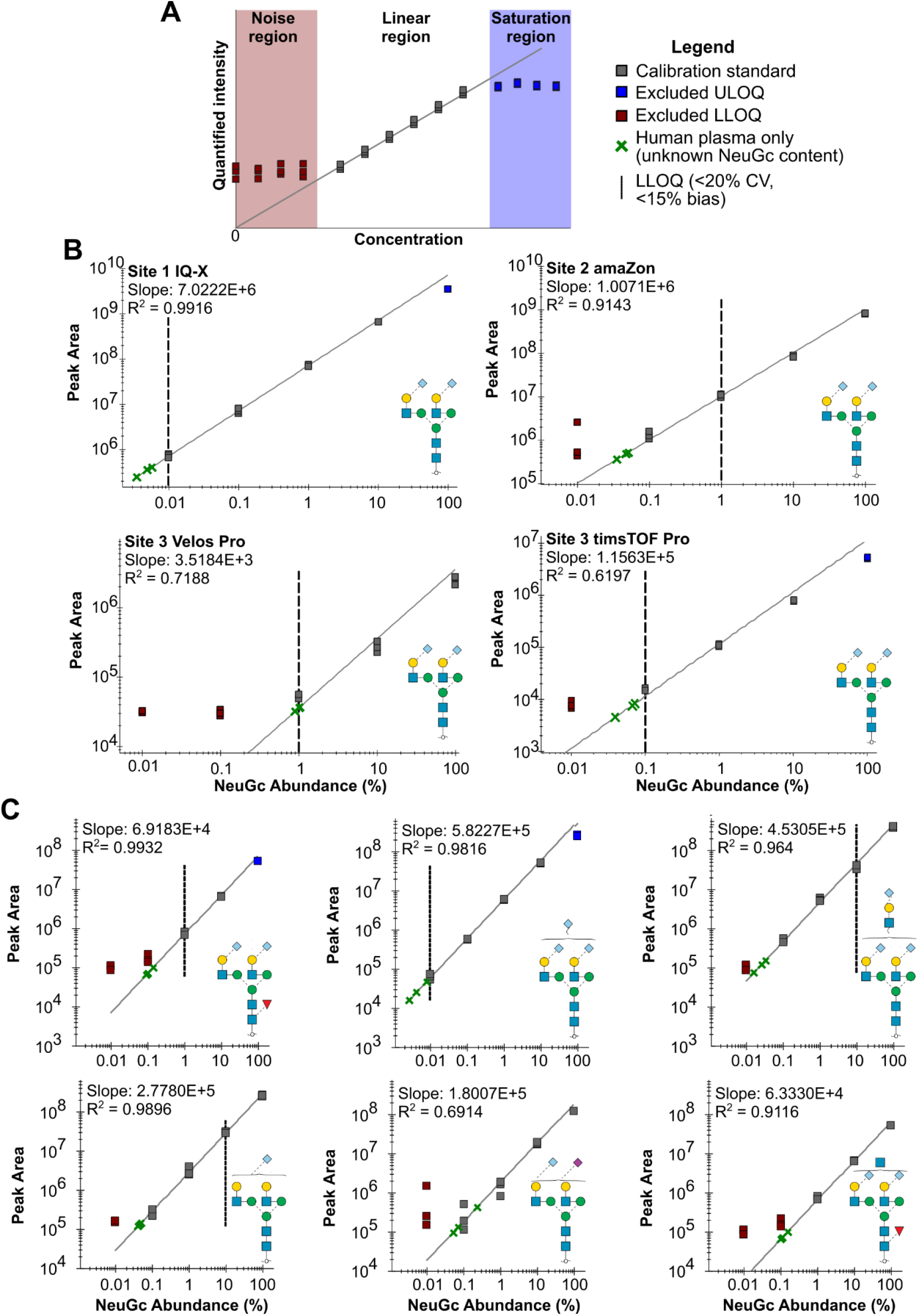
Quantitative assessment of Neu5Gc benchmark standards reveal instrument and structure-specific variation and accuracy. **A** Terminology guidance for a calibration curve. **B** Site comparisons for calibration curves of the most abundant Neu5Gc-bearing *N*-glycan. **C** Calibration curve comparisons for structures acquired at Site 1.

In the case of Site 1 (IQ-X, **Figure 3B**), the most concentrated sample saturated the MS, and was therefore outside the linear range, above the ULOQ, while the remaining concentrations were above the LLOQ. Comparing the most abundant *N*-glycan across the remaining sites, various reasons were responsible for the LLOQ being higher, such as variation at low concentrations (Site 2 amaZon), or loss of linearity (<15% bias, Site 3 Velos Pro). As expected, newer instruments leveraging more recent advancements (IQ-X and timsTOF Pro) resulted in a more sensitive LLOQ. Site 1 served as the benchmark for assessing dynamic range. By injecting the largest amount of material, it was the only site to detect Neu5Gc-bearing *N*-glycans in neat human serum, below the intensity of the lowest-concentration mixture, at up to 0.01% of total sialylation. At all other sites, the human sample was indistinguishable from noise.

Moving beyond the biantennary, disialylated *N*-glycan, other structures were quantified across the sites, each at a lower intensity and therefore higher LLOQ (**Figure 3C**). The second most intense Neu5Gc glycan, a trisialylated biantennary *N*-glycan, had a similar calibration curve to the most abundant glycan. For less intense glycans, loss of linearity and greater variation caused the LLOQ to be higher.

For the past decade, the gold-standard method for analysing sialic acids has been LC-FLR of fluorescently labelled monosaccharides, an example workflow of which is shown in **Figure 4A**. This approach has been widely adopted due to its selectivity for sialylated monosaccharides (Bardor, Nguyen et al. 2005, Martin, Muotri et al. 2005, Samraj, Läubli et al. 2014, Samraj, Läubli et al. 2014, Ye, Mu et al. 2020). By calibrating unknown sialic acid concentrations with a standard curve of known sialic acid concentrations, the absolute amount of an observed monosaccharide can be calculated based on the defined relationship between amount and fluorescence intensity, as demonstrated in **Figure 4B**.

**Figure 4.**
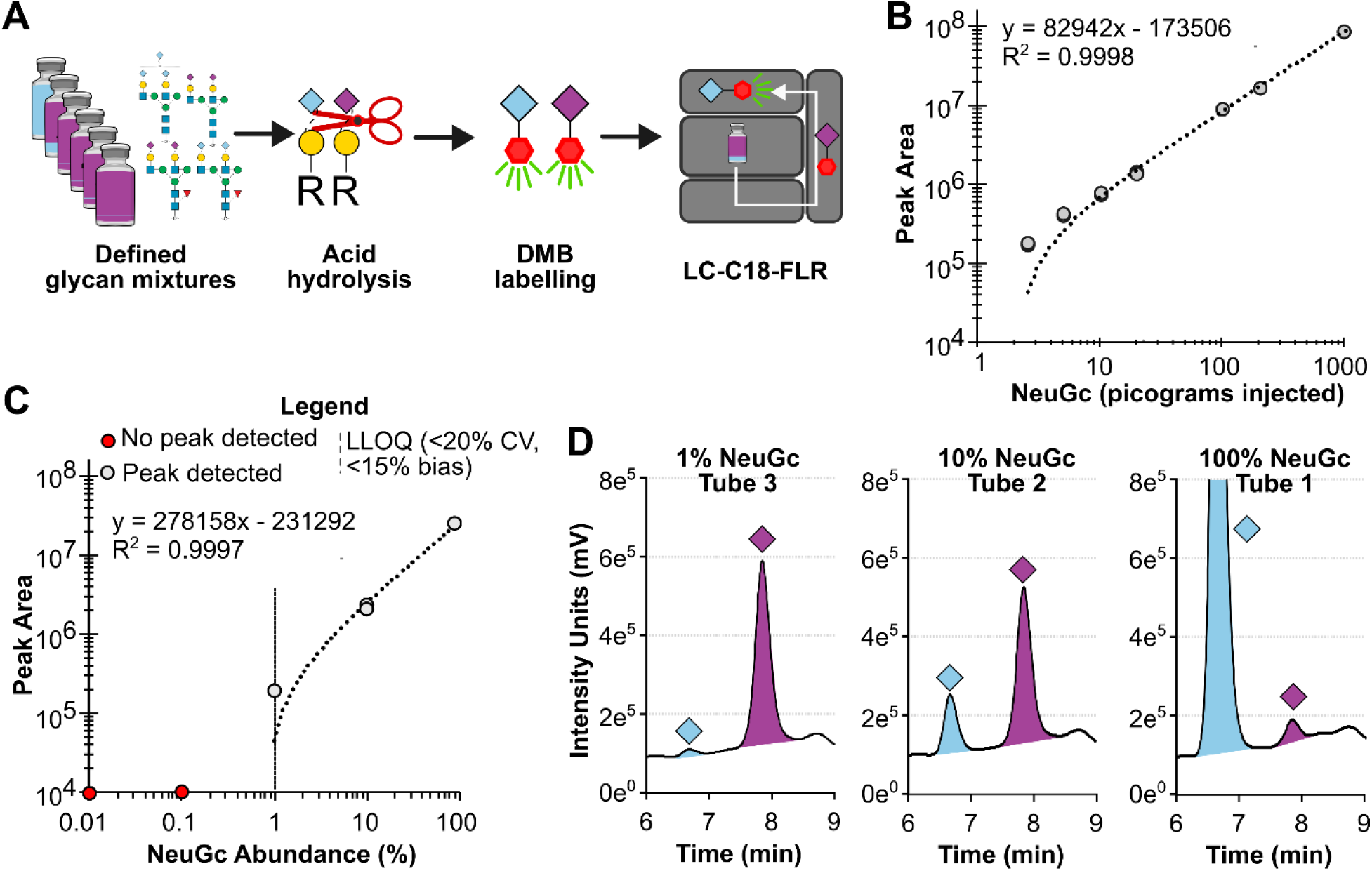
Methodology limitations for gold-standard monosaccharide quantitation drive Neu5Gc underestimation in human samples. **A** Monosaccharide analysis workflow for applied to Neu5Gc benchmark standards. **B** Monosaccharide standard calibration curve reveals linearity across two orders of magnitude. **C** Two orders of magnitude dynamic range are insufficient to detect Neu5Gc in human serum. **D** Example chromatogram of a Neu5Gc and Neu5Ac mixture at the limit of detection.

When this method was applied to the benchmark Neu5Gc sample set previously characterised by LC-MS, a peak corresponding to Neu5Gc was detected in the three most abundant samples (**Figure 4C**). In contrast to LC-MS, where signal was detected but variable beyond 20% co-efficient of variation at most sites, the Neu5Gc peak for the remaining three samples could not be confidently detected, owing to the consistent background noise inherent to the assay. The monosaccharide analysis results also reflect the limitations of the assay with the 1:100 dilution factor sample barely registering a Neu5Gc peak (**Figure 4D**). Importantly, this approach also provided absolute concentrations of both Neu5Ac and Neu5Gc monosaccharides across the benchmark samples (**Supplementary Table 1**), contextualising the concentration ranges that defined the sensitivity and quantitative limits of both LC-MS and LC-FLR assays.

## Discussion

### Analytical platforms vary in their Neu5Gc detection capabilities

The multi-site benchmarking study revealed clear LC-MS-specific differences in the LLOQ for Neu5Gc containing *N*-glycans, emphasizing that the conclusions drawn regarding the presence of Neu5Gc in human samples are inherently platform-specific. Only Site 1 was able to detect Neu5Gc-bearing *N*-glycans in neat human serum, at ∼0.01% total sialylation. On the other platforms employed in this work, this level was consistent with noise. Each site injected a different amount of *N*-glycans, however, with exception to the timsTOF, the differences in dynamic range exceeded what the 10–100× differences in injected material could account for.

Idotp values deteriorated with increasing dilution, confirming that MS1 isotopic distribution is a reliable proxy for signal confidence at low abundance and should be used as a routine quality metric in Neu5Gc detection workflows. This approach does have its limitations as at one site, significant carryover was observed, which emphasises the importance of block randomization and/or re-injection after blank acquisitions to limit the effect of carryover on false detections. Idotp measurement itself also has limitations, and cannot discriminate isomeric compositions, so orthogonal methods and metrics (e.g. MS2 and fragmentation coverage) are needed to resolve these interferences.

Fragmentation of *N*-glycans was, as expected, also platform-specific, with resonant CID instruments (amaZon, Velos Pro) failing to generate the diagnostic monosaccharide product ion for Neu5Gc, 306.08 *m/z*, which was observed on beam-type instruments (timsTOF Pro, IQ-X). Despite this, the neutral and charge losses of Neu5Gc were observed across all platforms, as well as larger product ions containing Neu5Gc (*e*.*g*. 671.22 *m/z {[M-H]*^*-*^*}*). In the untargeted methods used here, fragmentation of *N*-glycans of interest is not guaranteed, but the use of targeted methods (e.g. PRM/tMS2) may offer a solution for instruments where noise overwhelmed signal, given the specificity of precursor and product-ion targeting.

### The limitations of gold-standard LC assay for sialic acid content

Comparison between LC-MS and the current gold-standard DMB-LC-FLR approach (Manzi, Diaz et al. 1990, Martín, Vázquez et al. 2007) demonstrated that released monosaccharide analysis could only detect Neu5Gc across two orders of magnitude, whereas LC-MS extended detection to three or more orders of magnitude despite measuring each Neu5Gc-containing structure individually. This suggests that LC-MS offers superior specificity and sensitivity for Neu5Gc at the cost of more expensive equipment and more complex data analysis.

The two-order of magnitude dynamic range of the DMB-LC-FLR approach proved insufficient to detect Neu5Gc at concentrations representative of human serum, contextualising why these assays have largely been unable to detect Neu5Gc in clinical human samples. The absolute quantitation afforded by the DMB standard curve was valuable in this context as it provided the real concentration of the monosaccharides, unattained by the LC-MS approach used here. A hybrid strategy combining the two, such as LC-MS of released monosaccharides, may be a natural next step for subsequent analysis. In principle, this hybrid technique could yield absolute concentrations (using stable-isotope-labelled monosaccharide standards) while retaining the specificity provided by MS.

### Opportunities for Neu5Gc as a biomarker

Detection of Neu5Gc in human serum at only one site, at trace levels (∼0.01%) supports the hypothesis that Neu5Gc has been systematically under-detected in human glycomic datasets. This likely can be attributed to a combination of inadequate instrument sensitivity and overly restrictive search spaces, where Neu5Gc can be left out of data analysis workflows due to isomeric complexity and a subsequent increase in false positives(Klein, Carvalho et al. 2024, Bienes, Yokoi et al. 2025), and the absence of a rigorous LLOQ evaluation as performed here.

Future studies evaluating Neu5Gc in human samples (whether across populations or in sample types beyond sera) should prioritise pairing this low abundance glycan epitope with high-sensitivity platforms rather than the DMB-LC-FLR approach, which is fundamentally unable to specifically detect or quantify Neu5Gc at concentrations found in neat pooled human sera.

### Conclusion

This study applied a matrix-matched calibration curve to released *N*-glycans for the first time, providing a reusable framework for benchmarking Neu5Gc detection across LC-MS platforms and sites. Within this framework, LC-MS resolved Neu5Gc-bearing *N*-glycans over three or more orders of magnitude and, at the site with the largest linear range, detected Neu5Gc in neat human serum at ∼0.01% of total sialylation, whereas the gold-standard DMB-LC-FLR assay was confined to two orders of magnitude and could not detect Neu5Gc at concentrations representative of human serum. These platform-specific limits strongly support that Neu5Gc has been systematically under-detected in human samples, and indicate that future studies should pair this low-abundance epitope with high-sensitivity LC-MS, ideally combining released-monosaccharide MS with isotopically labelled standards to recover absolute quantitation.

## Materials and methods

### Development of standard mixtures

All chemicals and reagents were purchased from Millipore Sigma (Castle Hill, Australia) unless specified otherwise. *N*-glycans were released from human serum (H4522-20ML, Sigma-Aldrich), and a proprietary Neu5Gc-rich glycoprotein mixture (Protea Glycosciences), as described in SUGA (Ashwood, Voelcker et al. 2025) with minor modifications. Briefly, approximately 1 mg of each protein mixture had cysteines reduced and alkylated, followed by protein precipitation by acidified methanol on DNA miniprep columns (Bioneer) (Mousseau, Pierre et al. 2023). After precipitation, protein was resuspended in 240 µL of 100 mM triethylammonium bicarbonate with 200 units of PNGase F (Promega) and held at 37 °C for 18 hours. This preparation was repeated in quintuplicate and in duplicate for the human serum and Neu5Gc rich material, respectively.

Following *N*-glycan release, *N*-glycans were eluted with 400 µL of 0.1% formic acid and then immediately dried by centrifugal evaporation. Once dry, *N*-glycans were reduced and desalted as described (Jensen, Karlsson et al. 2012, Moh, Dalal et al. 2024), with 1 M NaBH_4_ in 50 mM KOH at 50 °C for 3 hours. After reduction, *N*-glycans were desalted by Supelclean ENVI-Carb solid phase extraction (100 mg) with 400 µL acetonitrile/0.1% formic acid conditioning, 1.5 mL water/0.1% formic acid equilibration, sample loading, 1.5 mL water/0.1% formic acid desalting, and 400 µL 50:50 acetonitrile: water with 0.1% formic acid elution. Eluted, desalted glycans were then dried by centrifugal evaporation.

Each sample replicate was analysed by LC-MS, observed intensity of the most abundant glycan was recorded, pooled by sample identity, and diluted to be at the same approximately MS intensity across two final tube pools. *N*-glycan mixtures were made of the two components, with the Neu5Gc *N*-glycans serially diluted by Neu5Ac *N*-glycans 10-fold to form a total of 4 samples, for a total of 6 samples when including the Neu5Ac and Neu5Gc starting material as separate samples. Each tube contained an *N*-glycan equivalent of a release from 500 µg of protein. *N*-glycans were then dried and mailed at room temperature with receipt within 5 days. A total of 15 benchmark sample sets were generated within a single batch.

### LC-MS data acquisition methodologies

Initial characterisation by LC-MS was performed at Site 1. Glycans were separated with a Thermo Fisher Scientific Vanquish Horizon HPLC (San Jose, USA) and ionised into an Orbitrap IQ-X Tribrid mass spectrometer (San Jose, USA). A Thermo Fisher Hypercarb porous graphitised carbon (PGC) column (Lithuania, 100 mm length by 1 mm internal diameter, 3 µm particle size), held at 90 °C, was used for all separations. Mobile phase A composed of water and mobile phase B composed of acetone with 5 mM HFIP and 5 mM butylamine added.

Three sites (four different instruments) were sent sets of the benchmark standards with instructions (fully detailed in the Supplement). Briefly, glycans were supplied dry and the only variables between sites were the resuspension volume and buffer.

Each laboratory group’s specific approach is summarized in the Supplement, with chromatography and mass spectrometry equipment and parameters in Table S2. Overall, the following methodology was consistent: Reduced *N*-glycans were separated on a porous graphitised carbon column over a binary gradient with the eluent subsequently analysed online by a mass spectrometer operated in negative mode, obtaining both MS1 (precursor) and MS2 (product) spectra.

### LC-MS data analysis

Upon acquisition, raw files were sent to the corresponding authors for subsequent analysis in Skyline-daily, using an assay template developed by Site 1. Spectrum visualization was performed with Freestyle 1.8 SP2 (Thermo Fisher Scientific), FlexAnalysis 3.4 (Bruker), and DataAnalysis 6.2 (Bruker). LC-MS raw files were analysed using GlyCombo (v1.2) to assign glycan compositions to precursor *m/z* values, identify the most intense MS2 scans for each structure for annotation in GlycoWorkbench, and construct the Skyline assay detailing *N-*glycan compositions. For Site 1, mzML input was used with an error tolerance of 25 ppm, reducing end specified as free, derivatisation as native and adducts set as M-H^-^. Monosaccharide search space was: Hex 4-12, HexNAc 2-8, dHex 0-1, Neu5Ac 0-3, Neu5Gc 0-3.

Skyline-daily (v25.1) was used to integrate the first three isotopic peaks with mass analyser set to centroid at 15 ppm mass accuracy. These isotopic integrations were used to quality filter identifications (>=0.85 isotope dot product (idotp)) and quantify glycans. The idotp value of 0.85 was empirically selected to remove poor quality MS1-matches (caused by monoisotopic peak misassignment, incorrect charge assignment, and poor signal to noise ratios) while preserving high-quality matches.

### Monosaccharide analysis sample preparation

An equal volume of 4 M acetic acid (Merck) was added to each sample. Samples were then incubated at 80 °C for 3 hours to release sialic acids. Afterwards, 5 μL of each hydrolysed sample was added into 20 μL of 1,2-Diamino-4,5-methylenedioxybenzene dihydrochloride (DMB; Merck) labelling solution and then incubated at 50°C for 2.5 hours in the dark. DMB labelling solution contained Milli-Q water, 7.6% v/v glacial acetic acid, 0.74 M β-mercaptoethanol (Merck), 52 µM sodium hydrosulfite (Merck), and DMB at 1.6 mg/mL. Following this incubation, the DMB-labelled samples were diluted in 100 µL of ice-cold Milli-Q water. Samples were filtered using 0.22 μm syringe filters (Kinesis) and centrifuged at 900 × g for 5 min prior to analysis.

The DMB-labelled samples (Ex = 373 nm, Em = 448 nm) were then analysed on a Prominence-i HPLC (Shimadzu) using an XBridge Shield RP18 column (Waters) under isocratic elution in 86% Milli-Q water, 7% methanol, and 7% acetonitrile with chromatograms analysed with an intensity multiplier of 0.001. A Neu5Ac and Neu5Gc (Agilent) standard curve (Supplementary Figure 1) and a DMB labelling solution only control were included in analysis.

## Supplementary figures and data

1. **Supplementary Table 1** Absolute concentration of Neu5Gc and Neu5Ac monosaccharides across the 6 defined glycan mixtures as determined by DMB-LC-FLR analysis
2. **Supplementary Table 2** Total amount of material used per LC-MS injection
3. **Supplementary Figure 1** Standard calibration curve applied to calculate sialic acid concentration
4. **Supplementary Figure 2** Mobilogram for the most abundant sialylated *N*-glycans
5. **Supplementary Methods**

## Acknowledgements

This research was facilitated by access to Sydney Mass Spectrometry, a core research facility at the University of Sydney. C.A. is the director of Protea Glycosciences, a company that provides fee-for-service glycomics assays, software, and analytical standards, including the calibration samples described in this work. The other authors have no conflicts of interest to declare.

## Funding

This work was supported by the Ovarian Cancer Research Foundation to ND, CJD and MPJ; Australian Research Council to DK [DP240100832, DP250104387]; and the National Health and Medical Research Council [GNT2018947 to DK, GNT2026356 to MPJ].

### Abbreviations

CID: Collision-induced dissociation
CMAH: Cytidine monophosphate N-acetylneuraminic acid hydroxylase
DDA: Data-dependent acquisition
dHex: Deoxyhexose
DMB: 1,2-Diamino-4,5-methylenedioxybenzene
DMB-LC-FLR: 1,2-Diamino-4,5-methylenedioxybenzene–liquid chromatography–fluorescence detection
ESI: Electrospray ionization
Ex: Excitation wavelength
FLR: Fluorescence (detection)
HCD: Higher-energy collisional dissociation
HESI: Heated electrospray ionization
Hex: Hexose
HexNAc: *N*-acetylhexosamine
HFIP: Hexafluoroisopropanol (1,1,1,3,3,3-hexafluoro-2-propanol)
HPLC: High-performance liquid chromatography
ID: Internal diameter
idotp: Isotopic dot product
LC: Liquid chromatography
LC-MS: Liquid chromatography–mass spectrometry
LLOQ: Lower limit of quantitation
LOD: Limit of detection
[M-H]-: Deprotonated molecular ion
MALDI: Matrix-assisted laser desorption/ionization
MeCN: Acetonitrile
MMCC: Matrix-matched calibration curve
MS: Mass spectrometry
MS1: First-stage mass spectrometry (precursor ion scan)
MS2 / MS/MS: Tandem mass spectrometry (product ion scan)
*m/z*: Mass-to-charge ratio
Neu5Ac: *N*-acetylneuraminic acid
Neu5Gc: *N*-glycolylneuraminic acid
PGC: Porous graphitised carbon
PNGase F: Peptide-*N*-glycosidase F
SUGA: Swift Universal Glycan Acquisition
TEAB: Triethylammonium bicarbonate
tMS2: Targeted MS2 (targeted tandem mass spectrometry)
ULOQ: Upper limit of quantitation

## Data availability statement

Raw files are available on GlycoPOST (Watanabe, Aoki-Kinoshita et al. 2021) at https://glycopost.glycosmos.org/preview/15414602776a5429eb5a7fe with reviewer code 2328. Skyline assays and quantitation curves are available at Panorama (Sharma, Eckels et al. 2014): https://panoramaweb.org/NeuGcBenchmark.url.

