## Supplementary for "Development of a Matrix-Matched Calibration Curve for Multi-Site Quantification of Neu5Gc-Bearing N-Glycans"

Short running title: Matrix-Matched Calibration for Neu5Gc N-Glycan Assay

### Supplementary figures and data

1. **Supplementary Table 1** Absolute concentration of Neu5Gc and Neu5Ac monosaccharides across the 6 defined glycan mixtures as determined by DMB-LC-FLR analysis
2. **Supplementary Table 2** Total amount of material used per LC-MS injection
3. **Supplementary Figure 1** Standard calibration curve applied to calculate sialic acid concentration
4. **Supplementary Figure 2** Mobilogram for the most abundant sialylated *N*-glycans
5. **Supplementary Methods**

**Mailing address:** 73 The Avenue, Mount Saint Thomas, 2500, NSW, Australia

### Supplementary

**Supplementary Table I** Absolute concentration of Neu5Gc and Neu5Ac monosaccharides across the 6 defined glycan mixtures as determined by DMB-LC-FLR analysis

| Sample | Neu5Gc | Neu5Ac |
|---|---|---|
| **1** | 4.19 µg/mL (12.89 µmol) ±0.05 | 194.44 ng/mL (629.04 µmol) ±0.69 |
| **2** | 384.25 ng/mL (1.18 µmol) ±12.50 | 1.06 µg/mL (3.43 µmol) ±0.01 |
| **3** | 57.06 ng/mL (175.51 nmol) ±0.44 | 1.22 µg/mL (3.95 µmol) ±0.01 |
| **4** | Not Detected | 1.09 µg/mL (3.53 µmol) ±0.04 |
| **5** | Not Detected | 1.22 µg/mL (3.95 µmol) ±0.01 |
| **6** | Not Detected | 1.24 µg/mL (4.01 µmol) ±0.01 |

**Supplementary Table II** Total amount of material used per LC-MS injection

| Site (Instrument) | Tube reconstitution volume (µL) | Amount injected on column (µL) | Percentage used per injection |
|---|---|---|---|
| **1 (IQ-X)** | 66 | 10 | 15% |
| **2 (amaZon™)** | 500 | 6 | 1.2% |
| **3 (Velos)** | 500 | 3 | 0.6% |
| **3 (timsTOF Pro)** | 500 | 3 | 0.6% |

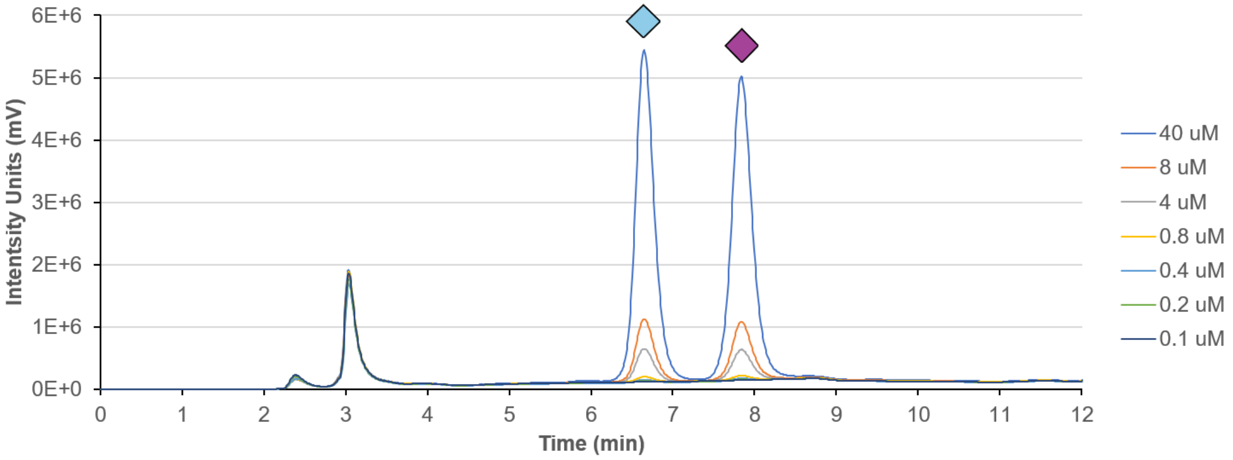


**Supplementary Figure 1** Standard calibration curve applied to calculate sialic acid concentration

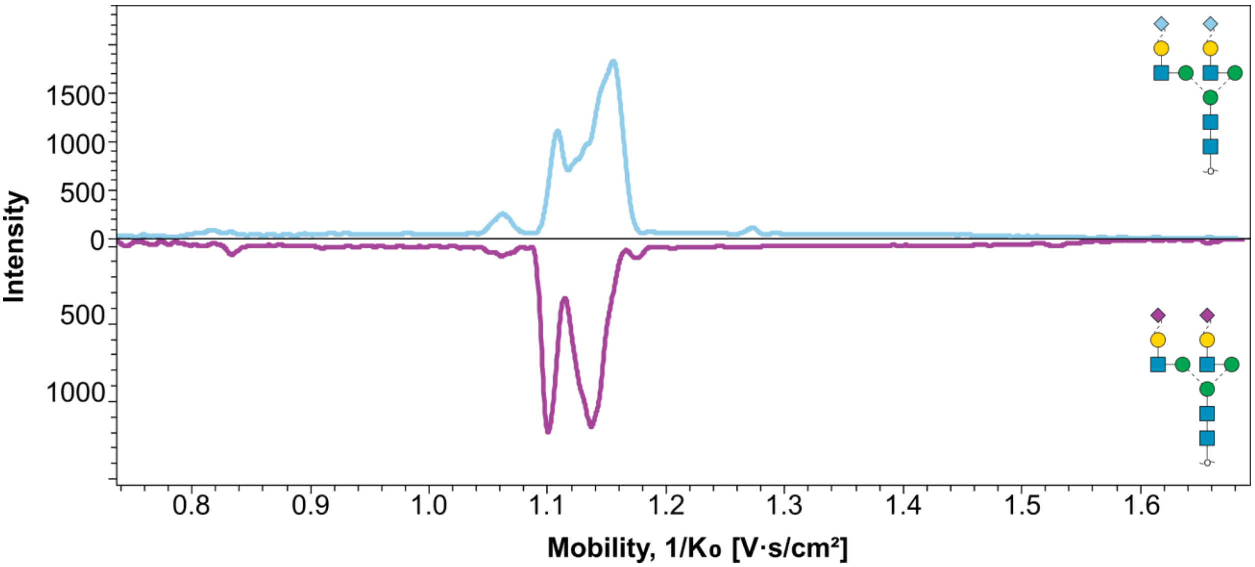


**Supplementary Figure 2** Mobilogram for the most abundant sialylated *N*-glycans

**Supplementary methods**

#### Site 1: IQ-X Method

Glycans were separated with a Thermo Fisher Scientific Vanquish Horizon HPLC (San Jose, USA) and ionised into an Orbitrap IQ-X Tribrid mass spectrometer (San Jose, USA).

### LC

A Thermo Fisher Hypercarb PGC column (Lithuania, 100 mm length by 1 mm internal diameter, 3 micron particle size), held at 90 °C, was used for all separations. Mobile phase A composed of water and mobile phase B composed of acetone with 5 mM HFIP and 5 mM butylamine added. LC separation was performed at 150 µL/min. Glycans were separated over a 10 min run, with 0-22% B over 4 min, 100% B held for 3 min, then 100% A for 2 min.

### MS

HESI spray voltage was set to -2.8kV, 30L/min of sheath gas and 20 L/min of auxiliary gas flow with no HESI temperature added. MS1 was operated as an Orbitrap scan from 570 - 1700 m/z at 240k resolution, with a maximum injection time of 502 ms and an AGC target of 1e6. MS2 operated in DDA mode with beam-type CID (HCD) fragmentation performed at 37% NCE with a 1.6 m/z isolation width. MS2 were acquired in the linear ion trap with a 1e5 AGC target and maximum injection time of 200 ms. Multiple precursors were subjected to MS2 until the target cycle time of 1.3 seconds was reached. Dynamic exclusion was enabled, excluding precursors for 6 seconds after MS2 events.

#### Site 2: amaZon method

### LC

Dried *N*-glycans were resuspended in water to a final concentration of ~ 1 µg/uL of equivalent protein and injection volumes were normalised to 6 µL. Samples were injected in technical triplicates and blank runs were monitored for sample cross contamination.
*N*-glycans were separated on a Hypercarb™ analytical column (30 mm x 1 mm ID, 3µm particle size, Thermo Fisher Scientific), maintained at 45°C, over a 65-minute method. Elution was performed using a binary mobile phase buffer system which consisted of 10 mM ammonium bicarbonate (Solvent A) and 70 % acetonitrile in 10 mM ammonium bicarbonate (Solvent B). *N*-glycans were loaded on the analytical column with 100 % Solvent A at a flow rate of 15 µL/min, followed by a 46-minute separation gradient initiated at 5.0 minute at a flow rate of 15 µL/min. *N*-glycans were eluted with increasing concentration of Solvent B, as follows: 1 – 14 % Solvent B over 1 minute, 14 – 25 % Solvent B over 19 minutes, 25 – 70 % Solvent B over 20 minutes, and 70 – 98 % Solvent B over 5 minutes. The column was re-equilibrated with 99 % Solvent A over 14 minutes prior to injection of the next sample.

### MS

Mass spectrometry acquisition was performed in negative mode. Initial scans were performed over a mass range of 460 - 1800 *m/z* at UltraScan mode with maximum accumulation time of 200 ms and an ion charge control target of 70,000. MS2 level fragmentation was obtained using collision induced dissociation activation over a mass range of 100 – 2200 *m/z*, with an isolation width of 4 *m/z* and fragmentation time of 28 ms. Preferred charge state for precursor selection was set to double and active exclusion was enabled after acquisition of 2 spectra and released after 1 minute.

**Site 3 timsTOF Pro method**

**LC**

An Agilent 1260 Infinity Capillary Pump was used for LC separation of a 30 mm x 1 mm ID porous graphitised carbon column with 3 µm particle size (Thermo Hypercarb), using 100% H2O, 10 mM AmBIC as mobile phase A and 70% MeCN, 10 mM AmBIC as mobile phase B. Glycans were separated at 20 µL/min with the following gradient profile: 0% B for 2 minutes, then ramped to 14% B for 1 minute, followed by a linear gradient to 30% B over 12 minutes, then another to 60% B over 8 minutes, followed by a hold at 100% B for 6 minutes. A 10 min post run at 0% B was used to re-equilibrate the column for the next injection. Column oven set to 50 °C.

**MS**

ESI spray voltage set to -4kV, 6L/min of dry nitrogen gas at 300 °C. MS1 and MS2 scan range: 100 – 2000 m/z, mobility range 0.7 – 1.6 V.s/cm$^2$, with 200 ms acquisition time. ICC and denoising was inactive. MS/MS operated in PASEF mode with beam-type CID fragmentation performed 75 eV with a 3 m/z isolation width. The top 6 precursors above 340 m/z with an intensity threshold of 5e3 were subjected to MS2 for a target cycle time of 2.68 seconds, with an active exclusion of 6 seconds.

**Site 3 Velos method**

**LC**

An Agilent 1260 Infinity Capillary Pump was used for LC separation on a 30 mm x 1 mm ID porous graphitised carbon column with 3 µm particle size (Thermo Hypercarb), using 100% H2O, 10 mM AmBic as mobile phase A and 70% MeCN, 10 mM AmBic as mobile phase B. Glycans were separated at 20 µL/min with the following gradient profile: 0% B for 3 minutes, then ramped to 14% B for 1 minute, followed by a linear gradient to 40% B over 36 minutes, then another to 56% B over 8 minutes, followed by a hold at 100% B for 4 minutes with a column equilibration at 0% B for 4 min. The column oven was set to 50 °C.

**MS**

MS was performed on a Velos Pro linear ion trap mass spectrometer. HESI spray voltage was set to -2.75kV, 13L/min of sheath gas and 7 L/min of auxiliary gas flow at 55 °C. MS1 was operated as a zoom scan from 400 - 2000 *m/z* and MS2 was performed as a normal resolution scan with the automatic scan range assuming a doubly charged precursor. MS/MS was operated in Top 5 DDA mode with resonant CID fragmentation performed at 33% NCE with a 1.5 m/z isolation width. MS1 AGC target was 3e4 with 10 ms maximum injection time. MS2 AGC target was 1e4 with 250 ms maximum injection time. Dynamic exclusion was enabled, excluding precursors for 15 seconds after MS2 events.
